# Physiological and pathological motor unit phenotypes coexist within single motor pools after cervical SCI

**DOI:** 10.64898/2026.06.04.730075

**Authors:** Devon R. Rohlf, Raul Sîmpetru, Dominik I. Braun, Daniela Souza de Oliveira, Matthias Ponfick, Alessandro Del Vecchio

## Abstract

Individuals with chronic cervical spinal cord injury (SCI) retain voluntary motor unit (MU) activation below the level of the lesion despite profound impairments in muscle relaxation, yet the MU-level mechanisms underlying this dissociation remain unclear.

Here, we investigated MU recruitment and derecruitment dynamics using high-density surface EMG (HD-sEMG) and intramuscular EMG (iEMG), in functionally paralyzed muscles across open- and closed-loop tasks in four participants with chronic cervical SCI. Building on prior work that distinguished task-modulated from non-modulated MUs, we combined closed-loop MU feedback with targeted intramuscular implants to determine whether tonic units remain accessible to voluntary control and whether distinct firing phenotypes coexist within individual motor pools.

Participants voluntarily recruited MUs and modulated discharge rates (45.2%), but a significant fraction (54.8%) of active MUs could not be derecruited during attempted relaxation. Quantitative analysis of 409 MUs revealed that derecruitment impairment spans a graded continuum rather than the binary distinction between modulated and non-modulated units reported previously. Three phenotypes, controllable (45.2%), modulated-tonic (42.8%), and tonic (12%), partition this spectrum based on derecruitment timing, discharge regularity, and firing persistence and coexist within the same motor pools, including within single muscle compartments sampled intramuscularly. Kaplan-Meier survival analysis confirmed graded derecruitment dynamics: controllable units ceased firing within 500 ms of rest onset, whereas most tonic units persisted throughout the observation window (log-rank *χ^2^*= 164.10, *P* < 0.001). Tonic units displayed highly regular discharge consistent with intrinsic motoneuron excitation via persistent inward currents (PIC). Phenotypes were consistent across recording modalities and task contexts, and phenotype effects on derecruitment metrics exceeded movement-type effects by an order of magnitude.

These findings identify impaired MU derecruitment as a core feature of spastic paralysis, driven by maladaptive motoneuron and spinal network properties that preserved descending drive cannot fully counteract. A proof-of-concept spike-train-level temporal filter selectively suppressed tonic firing while preserving voluntary modulation (>94% correlation retained), demonstrating a physiologically grounded strategy for improving neural interfacing after SCI.

## Introduction

After spinal cord injury (SCI), motoneurons below the lesion can begin to exhibit tonic firing that persists in the absence of voluntary drive, generating sustained involuntary muscle contractions.^1,2^ This pathological involuntary activation of motor units (MUs) is a core feature of what is clinically termed spasticity. While the classic definition of spasticity emphasizes a velocity-dependent increase in tonic stretch reflexes,^7^ the broader clinical syndrome encompasses involuntary muscle activity, exaggerated reflexes, and impaired voluntary muscle relaxation.^3–5^ In this study, we operationally define spasticity as the pathological involuntary activation of MUs that interferes with voluntary motor control. Individuals with spasticity can often initiate movements but struggle to terminate them, leading to sustained muscle activation during attempted rest.^2,6^ Although spasticity is frequently attributed to reflex hyperexcitability,^7,8^ this does not fully explain persistent tonic MU activity at rest or the dissociation between voluntary activation and relaxation. Whether these impairments arise from disrupted supraspinal drive, maladaptive spinal circuitry, or intrinsic motoneuron properties remains a central challenge, particularly given the limited studies examining MU behavior in paralyzed humans.

MUs translate descending neural commands into muscle force, and decades of research have demonstrated that MU recruitment and rate modulation can be voluntarily controlled.^9,10^ MU firing is governed by synaptic input and PICs, voltage-dependent calcium and sodium currents that amplify and sustain motoneuron excitation.^11–14^ After SCI, loss of descending regulatory signals from the brainstem can lead to exaggerated PICs, enabling motoneurons to continue firing after voluntary drive has ceased.^1,15,16^ Such intrinsic excitation provides a plausible mechanism for sustained muscle activity, yet its impact on voluntary MU derecruitment has not been systematically examined.^17^ Whether such post-injury changes act uniformly across the motor pool or differentially across its constituent motoneurons is unresolved, yet the distinction is mechanistically central: uniform alteration implies global dysfunction of the pool, whereas differential alteration would imply a mosaic of preserved and impaired motoneurons that residual descending drive could still partially engage.

Several fundamental questions remain unresolved: to what extent is voluntary MU recruitment preserved in motor complete cervical SCI, and does impaired relaxation reflect failure of activation or disengagement?^2,18^ Do all MUs in agonist muscles exhibit pathological firing, or do normal and dysfunctional units coexist? Can task context modulate pathological firing? These questions are essential for understanding spastic pathophysiology and for interpreting neural signals in applications that decode voluntary intent from MU activity.^19–22^

Here, we investigated MU recruitment and derecruitment dynamics using high-density surface electromyography (HD-sEMG) and targeted intramuscular EMG (iEMG) across open- and closed-loop tasks. We hypothesized that derecruitment impairment in SCI is heterogeneous, reflecting graded disruption of the balance between residual inhibitory control and intrinsic motoneuron excitability, rather than a uniform failure of voluntary drive. Participants voluntarily recruited MUs and modulated discharge rates, demonstrating preserved supraspinal command.^20^ However, many MUs could not be voluntarily derecruited despite shared descending commands. We identified three distinct phenotypes, controllable, modulated-tonic, and tonic, reflecting progressively impaired disengagement,^17^ consistent across recording modalities and supporting an intrinsic spinal origin.^12,14^ The coexistence of distinct MU behaviors within the same motor nucleus suggests that residual supraspinal drive is insufficient to overcome maladaptive spinal reorganization and altered intrinsic motoneuron properties after SCI. Finally, we demonstrate that phenotype-informed spike-level temporal filtering can selectively suppress tonic firing while preserving voluntary signals, enabling more robust spinal cord-computer interfaces for motor augmentation after SCI.

## Materials and methods

### Ethics and Patient Information

All procedures were approved by the ethics committee of Friedrich-Alexander-University Erlangen-Nuremberg (applications 21-150B, 22-138Bm, and 23-271B) in accordance with the Declaration of Helsinki. All participants provided written informed consent. Intramuscular electrode implantation was performed by a licensed physician guided by ultrasonographic imaging. Full exclusion criteria are provided in Supplementary Methods.

Four participants (P1–P4) with chronic cervical SCI (C4–6, American Spinal Injury Association Impairment Scale (AIS) A & B) were recorded across three experimental modalities: closed-loop sEMG with visual feedback, open-loop sEMG without feedback, and open-loop iEMG without feedback. P1 and P2 participated in all three experiments; P3 participated only in iEMG recordings; P4 participated only in open-loop sEMG. Each experimental modality was conducted on separate days. The open-loop sEMG and iEMG protocols were each completed in a single session; the closed-loop sEMG protocol was performed across multiple consecutive sessions (P1: 5 sessions; P2: 4 sessions). Recording modalities were separated by weeks to months, depending on participant availability and clinical scheduling. Three movements common across all sessions were analyzed: index flexion, two-finger pinch, and fist closure/opening.

Subsets of the sEMG recordings from P1, P2, and P4 have been reported previously in the context of real-time MU–computer interfacing,^23^ spinal cord–computer interface control,^20^ latent manifold analysis of spared motor dimensions,^21^ and EMG decoding framework validation.^22^ The present study applies new analyses: phenotype classification, survival analysis, and temporal filtering, to these recordings and additionally includes intramuscular EMG data from P1, P2, and P3 not reported elsewhere. (Supplemental Table 3)

Participant identifiers were harmonized across recording modalities for clarity. Demographic and clinical data are provided in Supplementary Table 1.

### Acquisition Hardware

All experiments used the Quattrocento bioelectrical amplifier (OT Bioelettronica, Torino, Italy). SEMG was sampled at 2,048 Hz (monopolar, hardware bandpass 10–4,400 Hz). IEMG was sampled at 10,240 Hz (bipolar, hardware bandpass 10–4,400 Hz). A ground reference was placed at the elbow. The skin was cleaned with an alcohol solution prior to electrode placement.

## Electrodes

Surface recordings used 64-channel high-density grids: 8×8 arrays (GR10MM0808, 10 mm interelectrode distance) and 13×5 arrays (GR08MM1305, 8 mm interelectrode distance; OT Bioelettronica). For intramuscular recordings, bipolar fine-wire electrodes (DIMW11050300102, Spes Medica, Genoa, Italy) were used, consisting of paired 50 μm stainless steel wires insulated with polytetrafluoroethylene (PTFE) and housed in a 23G cannula.

### Experimental Tasks

#### Open-Loop sEMG without Feedback

Participants P1, P2, and P4 were seated in their wheelchairs with the forearm of the most affected arm supported on the armrest. Three 8×8 and two 13×5 HD-sEMG grids were placed on the forearm. The 8×8 grids were positioned approximately one-third of the forearm length from the elbow, over the proximal muscle bellies of the flexor digitorum superficialis (FDS) and extensor digitorum (ED). The 13×5 grids were placed at approximately one-third of the forearm length proximal to the wrist over the flexor and extensor compartments.

A virtual hand interface was displayed on a monitor positioned in front of the participant.^20,22,24^ Participants were instructed to mimic the movements of the virtual hand as closely as possible. Because the virtual hand trajectory was pre-programmed and independent of the participant’s EMG, no neuromuscular feedback was provided; however, participants received continuous visual instruction via the displayed movement and verbal cues from the experimenter. Movements consisted of sinusoidal flexion–extension trajectories (0.5 Hz; 2 s cycles), preceded by a 2 s rest period and repeated 20 times per recording (42 s total). EMG recordings were acquired using the manufacturer’s acquisition software, OTBPlus (OT Bioelettronica), and synchronized with virtual hand movement onset via a 5 V TTL trigger signal connected to the amplifier’s auxiliary input.

Three movements common across all sessions were analyzed: index flexion, two-finger pinch, and fist closure/opening.

#### Closed-Loop sEMG with Feedback

Closed-loop recordings assessed MU behavior under conditions of controlled descending drive, using the NeurOne real-time MU–computer interface as previously described.^23^ EMG data were streamed to the NeurOne decomposition system using OTBLite, the manufacturer’s real-time streaming software (OT Bioelettronica). Participants P1 and P2 were recorded across multiple consecutive sessions (P1: 5 sessions; P2: 4 sessions). As in the open-loop protocol, participants were seated in their wheelchairs with the most affected forearm supported. One 8×8 grid was placed over the flexor muscle belly and one over the extensor muscle belly, approximately one-third of the forearm length distal to the elbow.

Each session began with a one-minute sinusoidal warm-up task in which the participant followed a virtual hand display. This recording was decomposed offline using DEMUSE to generate a separation matrix, which was then loaded into the NeurOne online decomposition system to serve as a spatial filter for real-time MU identification. Using this filter, participants tracked trapezoidal target trajectories consisting of a 3 s ramp up, 5 s plateau, and 3 s ramp down. The visual feedback cursor represented the mean MU discharge rate relative to the participant’s maximum voluntary effort established during calibration, allowing participants to modulate their activity to match the target. Six ramps per recording were performed at target effort levels of 20% and 60% of maximum voluntary effort, separated by 10 s rest periods.

#### Open-Loop iEMG without Feedback

Participants P1, P2, and P3 performed open-loop iEMG tasks. Participants were seated in their wheelchair (P1, P3) or reclining in a hospital bed (P2), the latter to accommodate the longer preparation time required for intramuscular electrode implantation. Fine-wire electrodes were implanted in functionally paralyzed target muscles under real-time ultrasound guidance (ArtUs EXT-2H, 15-MHz linear transducer, L15-7H40-A5; Telemed, Vilnius, Lithuania). Targets included: abductor pollicis longus (APL) and flexor pollicis longus (FPL) for D1; FDS compartments for D2, D3, and D4/D5. Full details of the implantation protocol, including ultrasound-guided localization, electrode positioning, and verification procedures, are provided in the Supplementary Methods.

Maximum voluntary contractions were obtained for each digit to establish proportional control scaling, with offsets computed from resting baseline activity to account for the influence of tonic MU activity. Participants then mimicked virtual hand movements (flexion tasks for individual digits, pinch grips, and fist) in 30–60 s blocks, adjusted for individual fatigue levels. As in the open-loop sEMG protocol, the virtual hand trajectory was pre-programmed and independent of the participant’s EMG; participants received visual and verbal instruction only. Movement speed, hold duration, and inter-cycle rest were adapted to each participant’s ability.

## Signal Processing

SEMG signals were visually inspected and channels with poor electrode contact or excessive artifacts were excluded. The monopolar signals were bandpass filtered (20–500 Hz, 4th-order Butterworth) and a 50 Hz notch filter was applied to remove power line interference. IEMG signals were bandpass filtered (10–1,500 Hz, 4th-order Butterworth). All offline filtering was performed in MATLAB.

To characterize the phase relationship between each motor unit’s firing rate and the kinematic reference, spike trains were converted to continuous firing rate traces by convolution with a 1 s Hanning window. Instantaneous phase of each smoothed firing rate and the kinematic reference was extracted via the Hilbert transform after linear detrending; sample-wise phase differences were wrapped to [−π, π] and retained only where both analytic-signal amplitudes exceeded their 30th percentile. The resulting distributions were visualized as 36-bin polar histograms and summarized per unit using circular mean, circular standard deviation, and mean resultant length.

### Decomposition

SEMG signals were decomposed using the convolution kernel compensation (CKC) algorithm^25^ implemented in the DEMUSE tool (v4.5; University of Maribor, Slovenia). Monopolar bandpass-filtered signals (20–500 Hz) were decomposed with 55 iterations per recording; no spatial filter was applied and 95% of electrode channels with the highest signal quality were selected for decomposition. Identified MU spike trains were visually inspected and manually edited by an experienced investigator following established procedures.^26^ MUs were accepted if the pulse-to-noise ratio (PNR) exceeded 30 dB and the silhouette score exceeded 0.9 after editing. Units with unstable or physiologically implausible discharge patterns were excluded.

IEMG was decomposed via manual spike sorting in EMGLAB.^27^ Motor unit action potential (MUAP) templates were identified by amplitude discrimination and waveform morphology. Individual spikes were assigned to templates based on shape matching, and assignments were manually reviewed for consistency across the full recording duration. Spike trains were inspected to confirm stable MUAP waveforms and physiologically plausible discharge rates.

### Motor Unit Phenotype Classification

MU derecruitment behavior after SCI spans a continuum from normal voluntary disengagement to complete failure of firing cessation. Rather than reflecting discrete pathological states, the degree of derecruitment impairment varies continuously across the motor pool, shaped by the balance between residual inhibitory control and intrinsic motoneuron excitability. To systematically characterize this heterogeneity, we defined three phenotypes that partition this continuum into operationally distinct categories based on converging derecruitment metrics. The controllable, modulated-tonic, and tonic labels designate regions along a graded spectrum of derecruitment impairment; individual MUs may occupy intermediate positions, and the modulated-tonic phenotype explicitly captures the transitional zone between preserved and failed voluntary disengagement.

MUs were classified using a four-criterion voting system based on derecruitment behavior. For each MU, the following metrics were computed: (1) coefficient of variation of inter-spike intervals (CV ISI) across the full recording (active and rest epochs combined), (2) fraction of rest epochs in which the unit was completely silent, (3) mean derecruitment delay (time from instructed rest onset to cessation of firing), and (4) per-epoch tonic firing fraction. Each criterion cast a vote of “controllable,” “tonic,” or “abstain” based on predefined thresholds (Supplementary Table 2). Threshold values were derived from prior literature on PIC-mediated tonic firing characteristics^1,12^ and applied without optimization to the present dataset. The voting system is designed to capture the same underlying phenomenon (sustained intrinsic motoneuron excitation) from multiple complementary angles: discharge regularity and firing rate characterize the real-time firing signature, while silent fraction and derecruitment delay capture epoch-level behavioral outcomes. This redundancy provides robustness across operational contexts, including real-time applications where epoch structure is unavailable.

A MU was classified as tonic if ≥3 criteria voted tonic, or ≥2 voted tonic with no controllable votes. Controllable classification required ≥3 controllable votes, or ≥2 controllable votes with no tonic votes. MUs not meeting either criterion were classified as modulated-tonic. Hard constraints were applied: units with derecruitment delay ≥500 ms could not be classified as controllable; units with delay ≤0.1 s could not be classified as tonic. Classification confidence was computed as the fraction of supporting criteria, reduced by 30% for units with fewer than three rest epochs and by 50% if CV ISI or silent fraction was undefined.

Because the phenotype boundaries are defined by physiological thresholds applied to derecruitment metrics, the resulting groups necessarily differ on those metrics. Group comparisons on classification inputs are therefore reported to quantify separation magnitude rather than as independent validation. The critical test is whether the phenotype partition captures meaningful structure beyond what alternative grouping variables explain: phenotype effects exceeded movement-type effects on the same metrics by an order of magnitude (see Results), and mean firing rate, which was not a classification input, also differed significantly across phenotypes (*ε*^*2*^ = 0.106, *P* < 0.001). Kaplan–Meier survival analysis confirmed graded derecruitment dynamics across phenotypes (log-rank *χ*^*2*^ = 164.10, P < 0.001), with controllable MUs derecruiting within 500 ms, modulated-tonic MUs declining gradually, and most tonic MUs persisting throughout the observation window. Phenotype distributions were consistent across all three recording modalities, confirming that the classification was not driven by decomposition artefact or recording technique.

Cross-session stability of phenotype assignments was assessed via MU tracking and is reported in the Supplementary Materials.

### Statistical Analysis

MU derecruitment metrics were compared across phenotypes (tonic, modulated-tonic, controllable) and recording modalities. Primary outcomes included derecruitment delay, fraction of silent epochs, CV ISI, and mean firing rate. Normality was assessed via Shapiro-Wilk tests; non-parametric methods were used due to significant departures from normality (all *P* < 0.001).

Phenotype comparisons used Kruskal-Wallis H-tests with Dunn’s post-hoc pairwise comparisons (Bonferroni-corrected *α* = 0.0167). Effect sizes were quantified using epsilon-squared (*ε*^*2*^), calculated as *H/(n−1)* where H is the Kruskal-Wallis statistic and n is total sample size.^28^ Following conventions for eta-squared in ANOVA,^29^ we interpret *ε*^*2*^ using thresholds of 0.01 (small), 0.06 (medium), and 0.14 (large), noting that no consensus exists for epsilon-squared interpretation specifically. Kaplan-Meier survival curves were constructed for derecruitment dynamics, with sustained firing beyond the observation window treated as right-censored; group differences were evaluated using log-rank tests.

### Spike-Train-Level Temporal Filtering

To demonstrate functional consequences of impaired derecruitment, we implemented a spike-train-level temporal filter to selectively exclude pathological firing while preserving voluntary activity. The filter operates on spike timing information from online MU decomposition.^20,23^

For each MU, firing behavior was evaluated in a sliding window of 15 consecutive inter-spike intervals (~2 s at typical firing rates). A segment was classified as pathological when mean firing rate fell between 7.5–8.5 Hz and windowed CV ISI was below 0.15, criteria derived from tonic phenotype characteristics and prior PIC literature. Spikes within detected pathological segments were excluded while spikes outside were retained, and filtered spike trains were smoothed using a 1 s Hanning window to yield a continuous control signal.

## Results

### Motor unit recruitment and derecruitment across recording modalities

Consistent with previous reports,^18^ participants with chronic cervical SCI and severe hand motor impairment voluntarily recruited MUs and modulated their discharge rates in accordance with instructed movement trajectories across all experimental modalities (Fig. 3). However, a substantial fraction of MUs failed to disengage during attempted relaxation (Fig. 3E–G), continuing to fire during rest periods even when participants were explicitly instructed to relax and when the same descending command was shared across the MU pool.

This derecruitment failure was not uniform. Across all recording modalities, 409 MUs were classified as controllable (45.2%, *n =* 185), modulated-tonic (42.8%, *n =* 175), or tonic (12%, *n =* 49) based on converging derecruitment metrics (Fig. 2). All three phenotypes were present in every participant (Supplementary Table 4), indicating that this heterogeneity arises within individual motor pools rather than from between-participant differences. Controllable MUs ceased firing rapidly during rest periods, often before the instructed rest phase, resulting in negative derecruitment delays and near-complete silence across rest epochs (Fig. 3A). Modulated-tonic MUs displayed intermediate behavior: they showed task-locked rate modulation during contraction phases but continued firing into rest before eventually ceasing activity after a delay (Fig. 3B). Tonic MUs exhibited sustained firing throughout rest periods, with prolonged positive derecruitment delays and no silent epochs (Fig. 3C). Intramuscular recordings confirmed sustained firing of individual MUs, revealing persistent single-unit activity in multiple muscles and digits (Fig. 4). Within the intramuscular recordings, which sample a single motor pool through compartment-targeted fine-wire electrodes, two or more phenotypes co-occurred within the same compartment in every participant studied with iEMG (P1, P2, P3), demonstrating that the heterogeneity is resolved to the level of a single motor pool.

**Figure 1:**
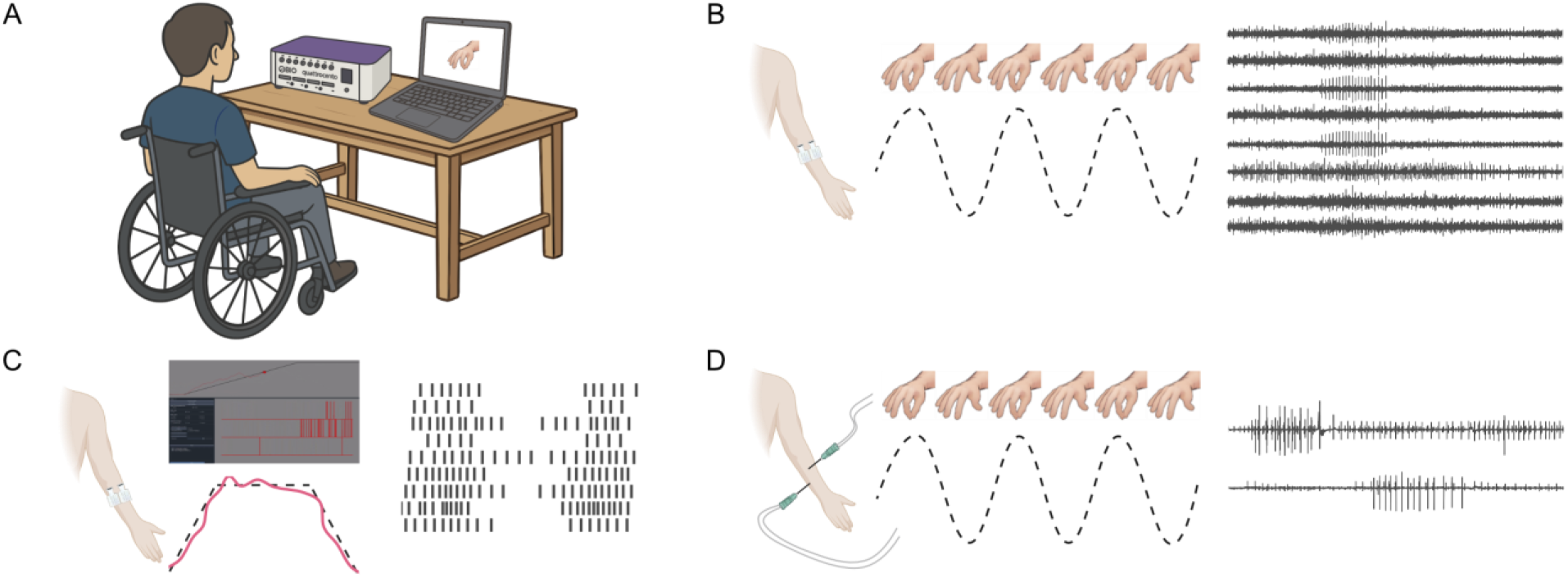
Experimental setup, recording modalities, and tasks. (A) Participants with chronic cervical spinal cord injury performed instructed hand/finger tasks from one of three experimental protocols while electromyography was recorded with a multichannel bioelectronic amplifier and task cues were presented via a virtual hand interface. (B) High-density surface EMG (HD-sEMG) open-loop task: participants attempted to follow a periodic sinusoidal flexion–extension trajectory displayed by the virtual hand without receiving feedback derived from their own EMG activity. HD-sEMG signals were recorded for offline analysis. (C) HD-sEMG closed-loop task: motor unit discharge rates extracted from online decomposition of HD-sEMG were used to provide real-time visual feedback, allowing participants to modulate activity to follow trapezoidal target trajectories. (D) Intramuscular EMG (iEMG) open-loop task: fine-wire electrodes recorded intramuscular activity during instructed movements without feedback, enabling single-MU resolution to confirm that sustained activity during rest is not a surface-signal mixing or decomposition artifact.

**Figure 2:**
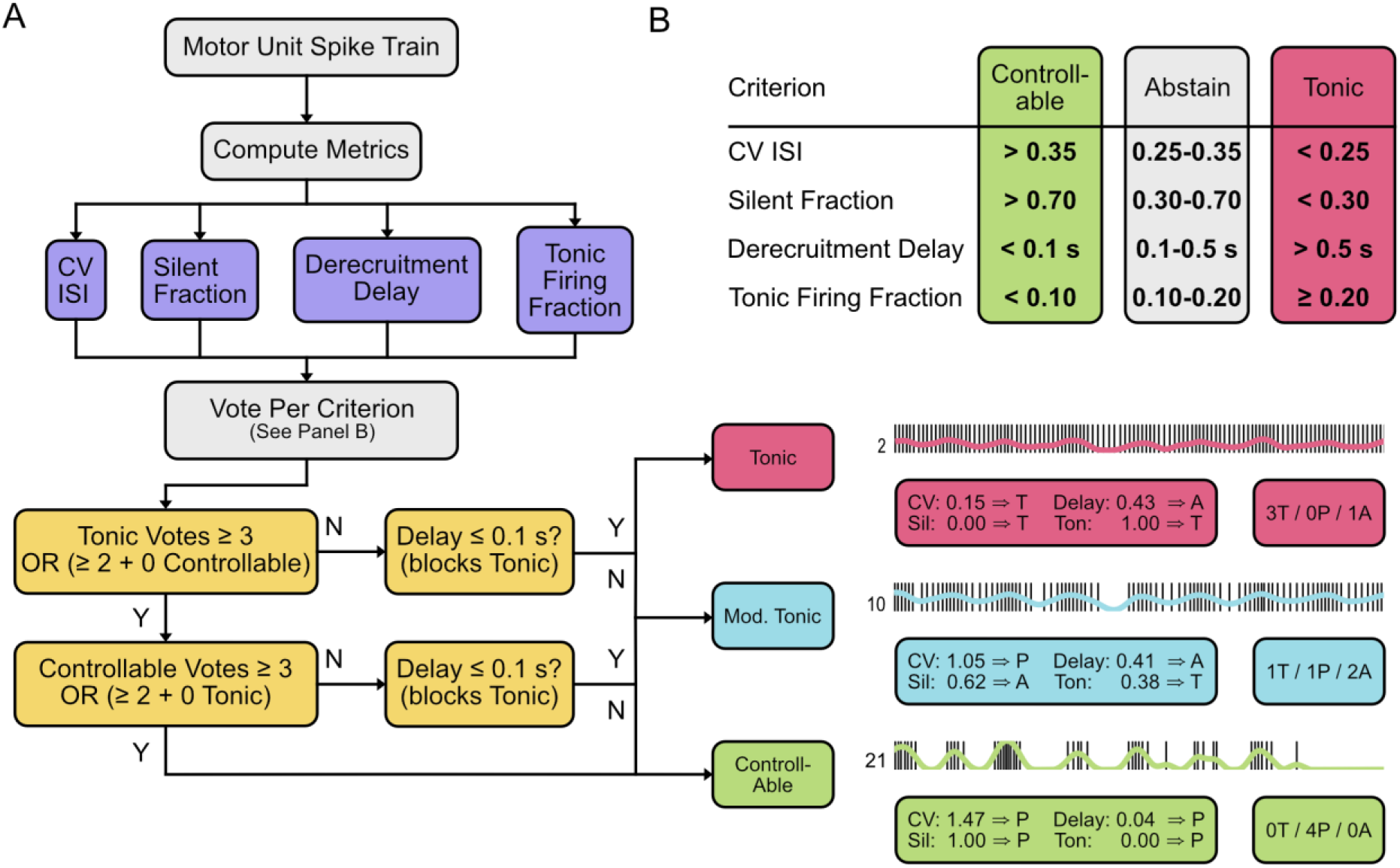
Voting-based motor unit phenotype classification system. (A) Decision flowchart for classifying motor units (MUs) into one of three phenotypes. Four metrics, shown in purple, are computed from each MU spike train: coefficient of variation of inter-spike intervals (CV ISI), silent epoch fraction, derecruitment delay, and tonic firing fraction. Each metric casts a controllable, tonic, or abstain vote according to the thresholds shown in Panel B. Votes are then evaluated through a decision cascade, shown in yellow: a supermajority, defined as ≥3 votes, or a simple majority, defined as ≥2 votes with no opposing votes, assigns the MU as tonic (red) or controllable (green). Hard physiological constraints override the vote count: derecruitment delay ≤0.1 s prevents tonic classification, whereas derecruitment delay ≥500 ms prevents controllable classification, redirecting the MU to the modulated-tonic phenotype (blue). MUs with mixed or insufficient evidence are likewise classified as modulated-tonic. (B) Threshold table for the four voting criteria. Green, grey, and red columns indicate value ranges that yield controllable, abstain, and tonic votes, respectively. Three representative MUs from a single open-loop recording illustrate the classification outcomes: MU 2, tonic with 3 tonic votes, 0 controllable votes, and 1 abstention; MU 10, modulated-tonic with 1 tonic vote, 1 controllable vote, and 2 abstentions; and MU 21, controllable with 0 tonic votes, 4 controllable votes, and 0 abstentions. Annotation boxes show each metric value and corresponding vote, and tally boxes summarize the vote counts.

**Figure 3:**
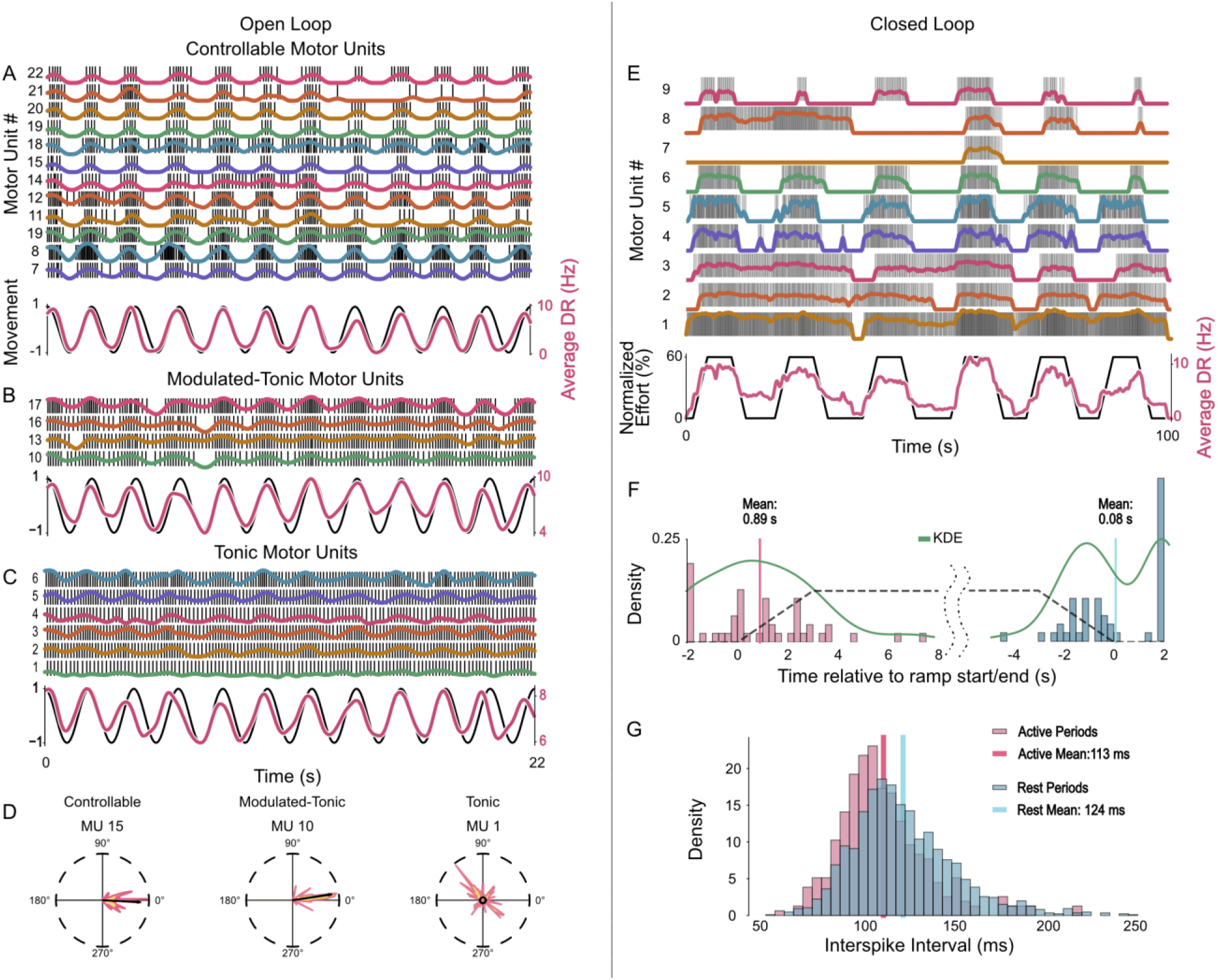
Preserved recruitment and impaired derecruitment of MUs during surface EMG task. (A) Controllable phenotype: motor units (MUs) are recruited and modulate discharge during active phases, then rapidly cease firing during intended rest, resulting in clear silent periods. (B) Modulated-tonic phenotype: MUs show task-locked modulation during effort but continue firing into rest before eventually derecruiting, indicating delayed disengagement. (C) Tonic phenotype: MUs remain active throughout intended rest with persistent, regular firing. Because phenotype classification reflects converging evidence across multiple derecruitment metrics rather than any single criterion, some tonic MUs retain partial discharge-rate modulation during active phases while still meeting tonic criteria through sustained firing across rest epochs, absence of silent periods, and low discharge variability. Tonic MUs occupy the most impaired end of the derecruitment spectrum but may retain partial modulation during voluntary effort. (D) Phase relationship between discharge-rate modulation and the movement trajectory for representative MUs, illustrating strong phase locking for controllable MUs and weak or absent coupling for tonic MUs. (E) Closed-loop recordings showing MU discharge during trapezoidal target tracking. Participants successfully modulated activity to follow the target, but tonic MUs continued firing during rest plateaus. (F) Kernel density estimates of derecruitment timing relative to movement offset, demonstrating delayed or absent derecruitment in modulated-tonic and tonic MUs. (G) Inter-spike interval distributions during active and rest epochs, highlighting increased discharge regularity during tonic firing.

**Figure 4:**
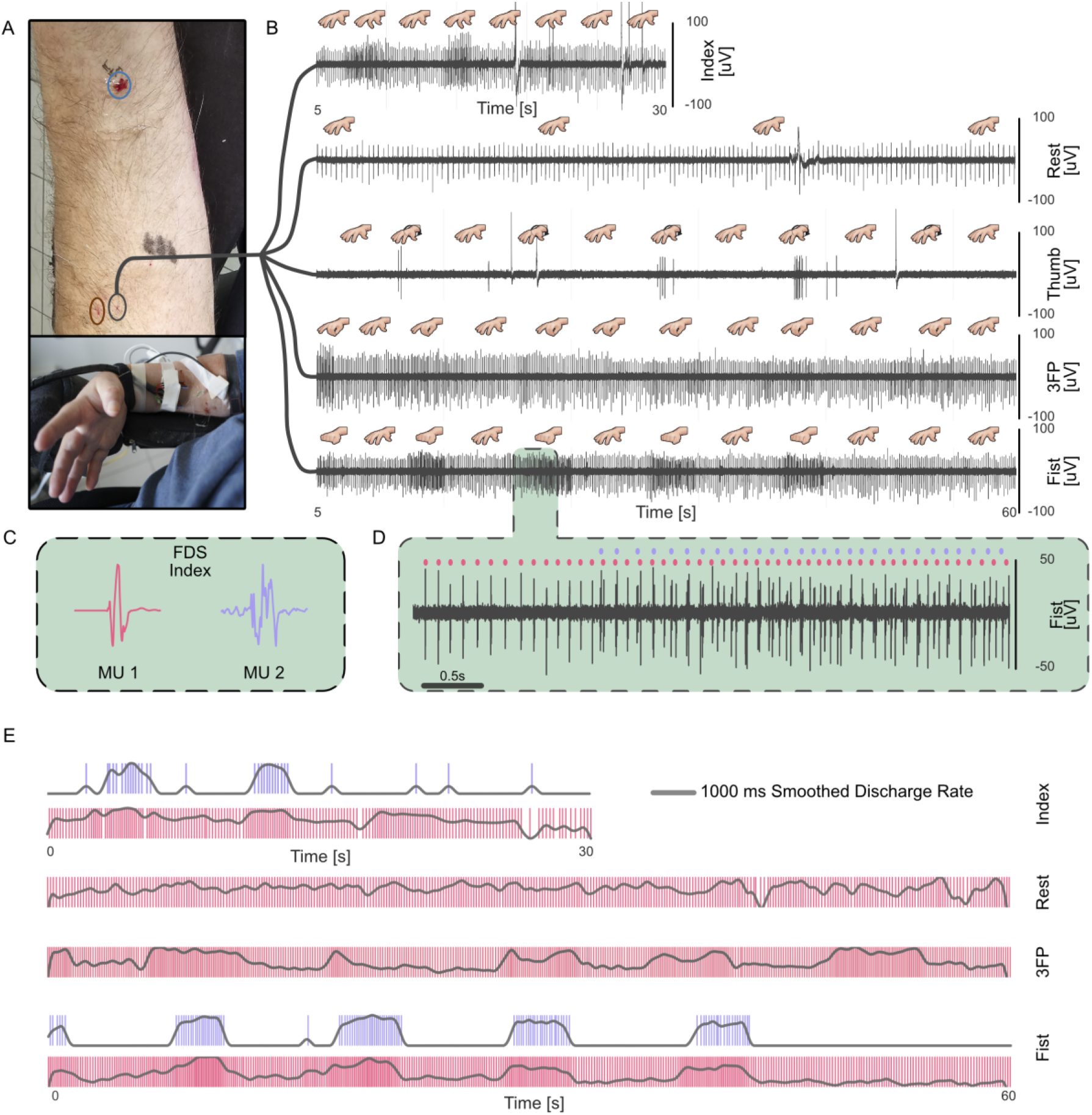
Intramuscular EMG confirms derecruitment failure at the single-motor-unit level. (A) Fine-wire implantation and recording configuration, with an example implant located in a digit-specific compartment of the flexor digitorum superficialis. (B) Representative intramuscular electromyography (iEMG) during attempted movements: sustained motor unit (MU) activity persists during intended rest across multiple task contexts recorded from the same implant. (C) MU action potential templates identified via manual spike sorting for two concurrently recorded units at the implant site. (D) Close-up view illustrating sustained firing, shown in red, interspersed with task-related modulated activity, shown in blue. (E) Smoothed discharge rates derived from intramuscular spike trains across task conditions, demonstrating preserved modulation during voluntary effort and persistent firing during rest. These recordings confirm that impaired derecruitment and tonic firing are present at the single-MU level and are not artifacts of surface electromyography signal mixing or decomposition.

### Time-to-derecruitment and discharge variability across phenotypes

Kaplan–Meier survival curves estimate the probability that a MU remains active (that is, has not yet derecruited) as a function of time after rest onset; a steeper decline indicates faster derecruitment, and units remaining active beyond the observation window are treated as right-censored. Because survival timing requires a precisely defined rest onset, this analysis used closed-loop recordings only, where the trapezoidal target specifies when relaxation should begin; the open-loop tasks lack a comparable kinematic reference. Time-to-event analysis revealed graded differences in derecruitment dynamics (Fig. 5A). Controllable MUs derecruited rapidly and completely, with all units ceasing firing within 500 ms of rest onset. In contrast, most tonic MUs remained active throughout the observation window (only 27.3% derecruited). Modulated-tonic units exhibited gradual decline, with 74.5% remaining active at 500 ms and 23.4% still firing at 1.5 s post-rest onset. Log-rank tests confirmed highly significant differences between survival curves (*χ*^*2*^ = 164.10, *P* < 0.001).

**Figure 5:**
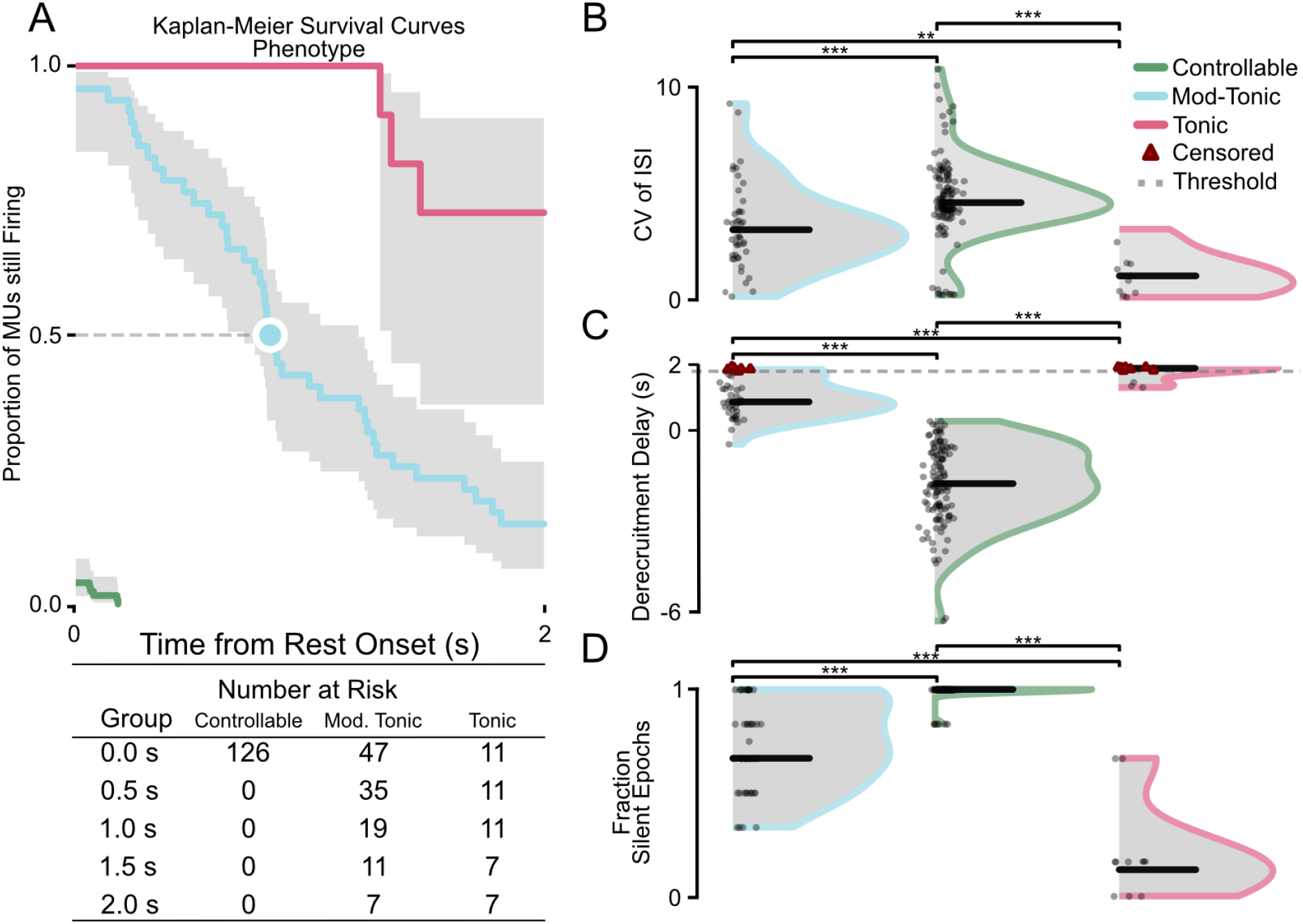
Phenotypes separate MUs by time-to-silence, discharge regularity, and rest suppression. (A) Kaplan–Meier survival curves, computed from closed-loop recordings where the trapezoidal target provides a defined rest onset, quantify the probability that a MU remains active after rest onset; sustained firing beyond the observation window is treated as right-censored. Controllable MUs derecruit rapidly, whereas tonic MUs often remain active throughout the observation window. (B) Discharge variability, quantified as the coefficient of variation (CV) of inter-spike intervals, differs across phenotypes, with tonic MUs exhibiting markedly reduced variability consistent with highly regular firing. (C) Derecruitment delay relative to movement offset captures graded impairment: controllable MUs cease firing near or before rest onset, modulated-tonic MUs show delayed cessation, and tonic MUs show the largest delays or non-derecruitment. MUs were treated as right-censored if they did not stop firing within 2 s of rest onset. (D) Fraction of silent rest epochs summarizes failure to cease firing during intended rest; controllable MUs are largely silent, whereas tonic MUs remain active. Statistical comparisons indicate significant phenotype effects (***P < 0.001).

Analysis of discharge timing revealed differences in firing regularity. Whole-recording CV ISI (computed across all inter-spike intervals from both active and rest epochs) was lowest for tonic MUs (median = 0.27), followed by modulated-tonic (1.45) and controllable MUs (4.20). Because this aggregate metric pools highly regular tonic segments with more variable voluntary segments, tonic units typically fell in the classification abstain range (0.25–0.35; Supplementary Table 2), and phenotype assignment depended primarily on the remaining three criteria. The low discharge variability of tonic MUs persisted across active and rest periods in both surface and intramuscular recordings.

### Comparison of phenotype and movement-type effects on derecruitment metrics

To quantify how much of the variance in derecruitment behavior the three-phenotype partition captures relative to other potential sources, we compared phenotype effects with movement-type effects on the same metrics (Fig. 5B–D). Derecruitment delay, fraction of silent epochs, and CV ISI differed significantly between phenotypes (all *P* < 0.001). Phenotype effects on derecruitment delay (*ε*^*2*^ = 0.746) were 13-fold larger than movement-type effects (*ε*^*2*^ = 0.056); similar ratios held for fraction of silent epochs (*ε*^*2*^ = 0.541 vs. 0.049) and CV ISI (*ε*^*2*^ = 0.431 vs. 0.056). Mean firing rate, which was not used as a classification input, also differed significantly across phenotypes (*ε*^*2*^ x = 0.106, *P* < 0.001), providing an independent confirmation that the partition reflects physiologically meaningful differences. Detailed statistical comparisons are provided in Supplementary Table 5.

### Spike-train-level temporal filtering of pathological motor unit activity

Spike-train-level temporal filtering was applied to selectively exclude pathological firing while preserving voluntary activity (Fig. 6). Using criteria derived from the tonic phenotype (7.5–8.5 Hz firing rate, CV ISI < 0.15; see Methods), the filter identified pathological firing segments within individual MUs. The same MU could exhibit excluded pathological firing during rest and preserved task-related firing during voluntary effort.

**Figure 6:**
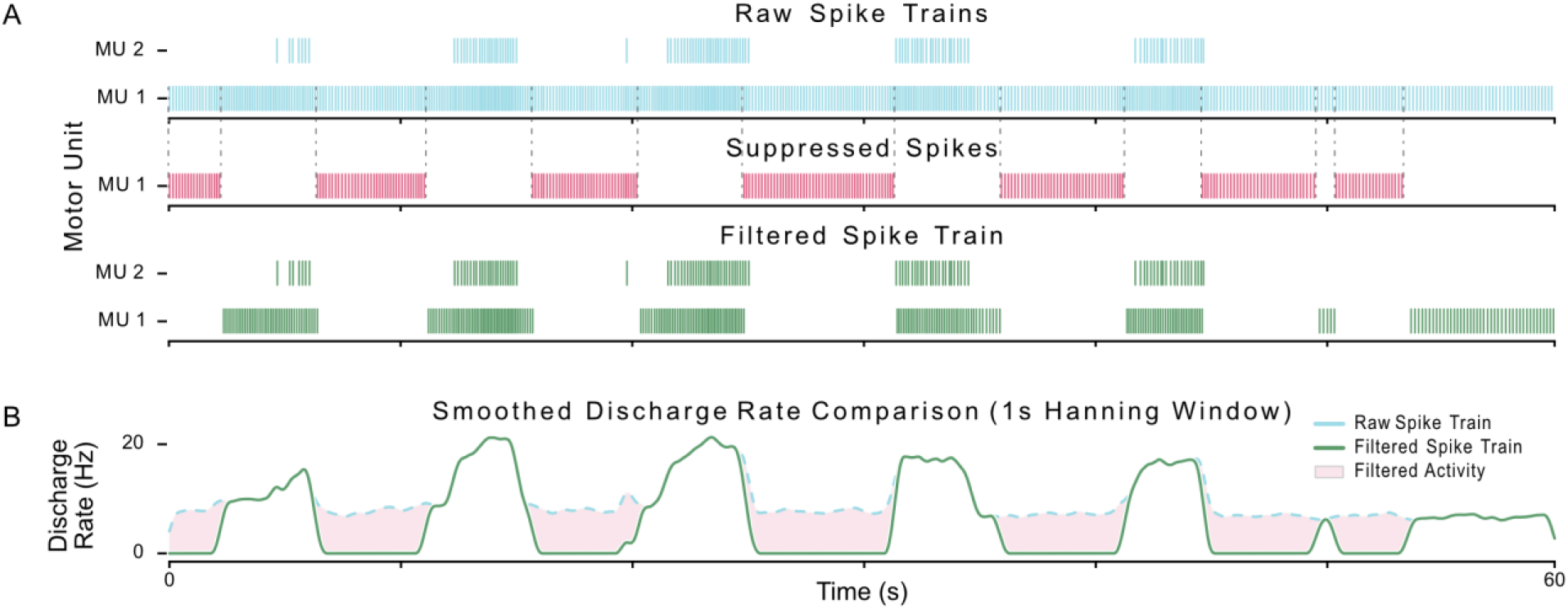
Spike-train-level temporal filtering suppresses tonic firing while preserving voluntary modulation. (A) Raw motor unit (MU) spike trains with tonic firing segments highlighted in red and voluntary task-related firing shown in blue. The same MU may exhibit both behaviors at different times. Filtered spike trains after removal of tonic activity are shown in green. (B) Smoothed discharge-rate signals, computed using a 1 s Hanning window from raw and filtered spike trains, show that filtering reduces tonic baseline activity while preserving the temporal structure of voluntary modulation. Shaded regions denote excluded tonic segments.

The smoothed discharge-rate signal from filtered spike trains remained highly correlated with the unfiltered signal during active task periods (*r* > 0.94). The filter selectively removed 43% of spikes from representative units, increasing the contrast between rest and activity periods. In units where tonic baseline activity was large relative to the voluntary modulation amplitude, filtering revealed task-locked discharge rate changes that were otherwise obscured in the unfiltered signal.

## Discussion

The central finding of this study is a dissociation in chronic motor complete cervical SCI: voluntary MU recruitment and discharge-rate modulation are preserved, but voluntary derecruitment is impaired across target muscles. Participants voluntarily recruited MUs and modulated their discharge rates across multiple recording modalities, demonstrating intact supraspinal access to the motor system.^19,20^ However, a substantial fraction of MUs failed to disengage during attempted relaxation, continuing to fire despite shared descending commands.^2,6^ This indicates that difficulty relaxing muscles in spastic paralysis reflects selective disruption of disengagement mechanisms rather than global failure of voluntary drive, constituting a defining feature of spastic MU behavior: the capacity to engage the motor system is retained, but the capacity to disengage it is selectively compromised.

This distinction is physiologically important. Recruitment reflects the capacity of descending pathways to activate motoneurons, whereas derecruitment requires withdrawal of excitation and engagement of inhibitory mechanisms. The preservation of recruitment argues against a primary cortical deficit and instead implicates maladaptive spinal mechanisms that selectively impair termination of MU activity.^18,30^

Derecruitment failure was not uniform across the MU pool. MUs segregated into three distinct phenotypes, controllable, modulated-tonic, and tonic, that coexist within the same muscles and individuals, indicating substantial intrinsic heterogeneity after SCI.^2^ Notably, even in uninjured individuals, a small fraction of MUs (~7%) are not modulated by task demands,^31^ suggesting the tonic phenotype may represent pathological amplification of a feature present within normal motor pools.

Previous work using HD-sEMG during open-loop attempted hand movements identified two broad MU populations after cervical SCI^18^: units whose discharge was modulated with the task and units that fired tonically or showed little task-related modulation. This raised two mechanistic questions. First, are tonic or weakly modulated units truly outside voluntary control, or can residual descending drive still modulate them under conditions of explicit neurofeedback? Second, do these units belong to the same motor pools as voluntarily recruited units, or do they reflect posture-related activity from other forearm muscles, a feature that can also occur in uninjured motor pools?^32^ The present study addressed these questions by combining closed-loop MU feedback with targeted intramuscular recordings from anatomically defined muscles. Together, these approaches show that derecruitment impairment is not a simple binary distinction between controllable and non-controllable MUs. Instead, MUs within the same motor pool span a continuum: controllable units can be recruited and derecruited closely according to task constraints, modulated-tonic units retain task-locked rate modulation but show delayed disengagement, and tonic units fail to derecruit during intended rest. Thus, the key pathological feature is not absence of voluntary recruitment, but impaired voluntary disengagement.

This framework refines the binary classification reported previously in overlapping participants, where MUs were categorized as task-modulated or non-modulated based on their phase relationship with reference kinematics.^18^ The comparable proportion of units with impaired derecruitment observed here, together with participant-level consistency across studies, suggests that both approaches capture the same underlying biological phenomenon. The present work extends that finding by resolving the broader task-modulated/non-modulated distinction into physiologically distinct derecruitment behaviors. In particular, modulated-tonic units may retain voluntary rate modulation during effort but still fail to disengage during rest, a distinction that becomes apparent only when derecruitment timing, discharge regularity, closed-loop control, and intramuscular motor-pool-level recordings are considered together.

Critically, the intramuscular recordings anatomically resolve this heterogeneity to the level of a single motor pool. In every participant studied with fine-wire electrodes, two or more phenotypes were recorded simultaneously from within the same muscle compartment, meaning that motoneurons innervating the same muscle, and presumably receiving common descending input during attempted movement, expressed qualitatively different derecruitment behaviors at the same time. The maladaptive process underlying spasticity therefore does not act uniformly across the motor pool. Some motoneurons retain near-normal disengagement, others fail to derecruit at all, and the population spans a graded continuum in between. The motor pool after chronic SCI is better understood as a mosaic of motoneurons in different post-injury states than as a uniformly altered population.

Controllable units exhibited rapid cessation of firing consistent with preserved inhibitory control. Tonic units continued firing throughout rest with no silent epochs, reflecting complete failure of voluntary disengagement. Modulated-tonic units showed delayed but ultimately successful derecruitment, suggesting that inhibitory mechanisms remain partially effective but are initially overwhelmed by sustained excitation, with intermediate discharge variability consistent with a mixed mode of synaptic and intrinsic excitation. Phenotype effects exceeded movement-type effects by more than an order of magnitude (e.g., *ε*^*2*^ = 0.746 vs. 0.056 for derecruitment delay), confirming that intrinsic MU properties rather than movement context are the dominant determinant of derecruitment behavior. This heterogeneity provides a physiological explanation for the mixed clinical presentation of spasticity.^6^ Cross-session MU tracking confirmed phenotype stability, with matched units showing 76% categorical agreement and significantly correlated continuous derecruitment scores.

The extremely low discharge variability of tonic units is inconsistent with synaptic drive alone and instead characteristic of sustained intrinsic depolarization from PICs. These amplify motoneuron excitation and exhibit hysteresis, allowing firing to persist after synaptic drive is withdrawn.^11,15,33^ These findings extend prior indirect evidence from paired MU PIC estimates and pharmacological studies by demonstrating, at the single-unit level, how intrinsic excitation manifests as derecruitment failure in humans with chronic SCI.

Intrinsic motoneuron properties alone, however, may not fully account for the observed phenotype heterogeneity. After SCI, the spinal cord undergoes spontaneous reorganization: spared corticospinal fibers sprout into denervated territories,^34–36^ and new propriospinal relay circuits form around the lesion site.^37^ In the absence of early targeted rehabilitation, this reorganization can be maladaptive, producing altered synaptic inputs to motoneurons^30^ and changes in network excitability that contribute to spasticity.^5,8^ This may explain why modulated-tonic and some tonic units retain partial rate modulation during active phases despite failing to derecruit: descending input reaches them through reorganized pathways that cannot support inhibitory termination. The interplay between maladaptive network reorganization and PIC-mediated intrinsic excitability could thus produce a continuum of derecruitment impairment, with the balance between these mechanisms varying across individual motoneurons. The within-pool heterogeneity observed intramuscularly implies that this reorganization is itself non-uniform. Differential changes in intrinsic excitability and PIC expression,^11,15^ heterogeneous sprouting of spared descending and propriospinal fibers^34–37^ onto motoneurons that previously received distinct input patterns, and uneven loss of segmental inhibitory control^5,8^ likely affect individual motoneurons within the pool to varying degrees rather than transforming the pool as a coherent unit. A direct consequence is that residual descending drive can still access and modulate the subset of motoneurons whose voluntary control has been spared, consistent with the partial preservation of voluntary recruitment we observed and with prior reports of preserved volitional MU activity below the level of the lesion.^18–20^ Earlier intervention through targeted rehabilitation or neuromodulation may guide plasticity before maladaptive networks consolidate,^38^ potentially reducing the proportion of motoneurons that develop tonic firing.

The coexistence of voluntary and pathological activity has direct functional consequences. Sustained tonic unit firing creates a mismatch between voluntary intent and motor output, complicating neural signal interpretation in interfaces that assume MU activity reflects movement intent. Our proof-of-concept temporal filtering demonstrates that derecruitment phenotypes can be exploited to distinguish voluntary from pathological activity at the spike-train level. This is particularly relevant for low-amplitude voluntary commands, where tonic firing from spastic MUs can generate large MU action potentials that dominate the composite signal and obscure movement intent. Such filtering could be integrated directly into existing real-time EMG decoding frameworks such as MyoGestic,^22^ which already supports online HD-EMG decomposition and proportional control in individuals with SCI. Phenotype-aware preprocessing would allow the decoder to operate on voluntarily modulated spike trains, potentially improving control fidelity without requiring retraining of the underlying model.^18^

These findings have important implications for contexts where MU activity serves as a proxy for movement intent. Distinct derecruitment phenotypes demonstrate that spastic paralysis involves a mixture of normally behaving, delayed, and persistently active MUs with pathological and voluntary firing potentially coexisting within the same unit at different times.^2^ This heterogeneity complicates EMG-based neural interfaces that have demonstrated preserved proportional control in SCI populations,^21,22^ as spastic firing patterns contaminate the decoded voluntary signal.

Common clinical approaches such as antagonist vibration or electrical stimulation act through global spinal mechanisms that broadly suppress motoneuron activity.^3,6,15^ While effective for reducing spasticity, these interventions may also attenuate voluntary MU firing, potentially eliminating signals required for neural interface control. Global suppression thus risks trading spasticity reduction for loss of usable control information.^19^

In contrast, spike-train-level temporal filtering selectively excludes pathological activity based on physiological firing characteristics without suppressing voluntary discharge. This emphasizes a fundamental distinction: clinical interventions aim to alter spinal excitability, whereas neural interfaces require accurate extraction of motor intent from available signals. Spasticity-related MU activity should not be treated as noise to be globally suppressed, but as a distinct physiological process that can be identified and selectively excluded.

Several limitations should be considered. The sample size was modest, and although derecruitment phenotypes were robust across participants, larger cohorts will be required to determine how phenotype prevalence depends on injury characteristics such as lesion level, chronicity, and spasticity severity. The small number of participants precludes formal hierarchical modeling of between-participant variance; however, all four participants individually exhibited all three phenotypes (Supplementary Table 4), and the phenotype proportions are consistent with what would be expected given differences in injury characteristics. The statistical analyses treat MUs as individual observations, which may overstate precision for population-level estimates of phenotype prevalence, but does not affect the within-participant finding that distinct derecruitment behaviors coexist within the same motor pools. In addition, PIC involvement was inferred from MU discharge characteristics rather than measured directly. While such features are well-established signatures of intrinsic motoneuron excitation, direct validation using paired MU techniques was not always feasible in the presence of tonic firing. Future studies incorporating paired MU analyses, pharmacological manipulation, or neuromodulatory interventions could provide more direct confirmation of the underlying mechanisms.

Longitudinal studies are also needed to assess the stability of derecruitment phenotypes over time and to determine whether MUs transition between states with training, neuromodulation, or recovery. Establishing the temporal dynamics of these phenotypes could enable their use as physiological biomarkers of disease progression or treatment response.

The spike-train-level filtering demonstrated here represents a proof-of-concept based on fixed, physiology-informed criteria. The current rate and regularity thresholds identify firing segments consistent with PIC-mediated activity, but equivalent firing patterns could in principle occur during sustained voluntary contractions in uninjured individuals. Incorporating task-context information, for example, restricting detection to epochs in which no voluntary activation is instructed, would disambiguate pathological from physiological tonic firing and improve specificity. Future work should explore such context-aware approaches, along with adaptive methods that account for non-stationary firing behavior.

## Conclusions

This study demonstrates that individuals with chronic cervical SCI retain the ability to voluntarily recruit MUs but exhibit a selective and heterogeneous impairment in voluntary derecruitment, the pathological involuntary MU activation that we argue defines spasticity at the MU level. Three distinct phenotypes, controllable, modulated-tonic, and tonic, capture this heterogeneity and are robust across surface and intramuscular recordings, open- and closed-loop tasks, and multiple participants. The highly regular discharge of tonic units implicates intrinsic motoneuron properties, consistent with PICs, rather than a global failure of supraspinal drive. Finally, spike-train-level temporal filtering demonstrates that pathological and voluntary firing can be separated within the same MUs, preserving voluntary control signals while selectively suppressing tonic activity. Together, these findings provide a mechanistic framework linking intrinsic motoneuron excitability to impaired muscle relaxation and offer a physiologically grounded strategy for improving neural signal interpretation after SCI.

## Data Availability

The data that support the findings of this study are available from the corresponding author, upon reasonable request.

## Acknowledgements

The authors thank all participants for their time and commitment to this research. We also extend our thanks to Daniel Haller and Patricia Bayer for their assistance in recording and curating data.

## References

1. Gorassini MA. Role of motoneurons in the generation of muscle spasms after spinal cord injury. Brain. 2004;127(10):2247–2258. doi:10.1093/brain/awh243

2. Zijdewind I, Thomas CK. Firing patterns of spontaneously active motor units in spinal cord-injured subjects. J Physiol. 2012;590(7):1683–1697. doi:10.1113/jphysiol.2011.220103

3. Adams MM, Hicks AL. Spasticity after spinal cord injury. Spinal Cord. 2005;43(10):577–586. doi:10.1038/sj.sc.3101757

4. Pandyan A, Gregoric M, Barnes M, et al. Spasticity: Clinical perceptions, neurological realities and meaningful measurement. Disabil Rehabil. 2005;27(1-2):2–6. doi:10.1080/09638280400014576

5. D’Amico JM, Condliffe EG, Martins KJB, Bennett DJ, Gorassini MA. Recovery of neuronal and network excitability after spinal cord injury and implications for spasticity. Front Integr Neurosci. 2014;8. doi:10.3389/fnint.2014.00036

6. Mottram CJ, Suresh NL, Heckman CJ, Gorassini MA, Rymer WZ. Origins of Abnormal Excitability in Biceps Brachii Motoneurons of Spastic-Paretic Stroke Survivors. J Neurophysiol. 2009;102(4):2026–2038. doi:10.1152/jn.00151.2009

7. Lance JW. The control of muscle tone, reflexes, and movement. Neurology. 1980;30(12):1303–1303. doi:10.1212/WNL.30.12.1303

8. Nielsen JB, Crone C, Hultborn H. The spinal pathophysiology of spasticity – from a basic science point of view. Acta Physiol. 2007;189(2):171–180. doi:10.1111/j.1748-1716.2006.01652.x

9. Basmajian JV. Control and Training of Individual Motor Units. Science. 1963;141(3579):440–441. doi:10.1126/science.141.3579.440

10. Heckman CJ, Enoka R. Motor Unit. Wiley; 2012. doi:10.1002/j.2040-4603.2012.tb00465.x

11. Bennett DJ, Li Y, Siu M. Plateau Potentials in Sacrocaudal Motoneurons of Chronic Spinal Rats, Recorded In Vitro. J Neurophysiol. 2001;86(4):1955–1971. doi:10.1152/jn.2001.86.4.1955

12. Li Y, Gorassini MA, Bennett DJ. Role of Persistent Sodium and Calcium Currents in Motoneuron Firing and Spasticity in Chronic Spinal Rats. J Neurophysiol. 2004;91(2):767–783. doi:10.1152/jn.00788.2003

13. Heckman CJ, Gorassini MA, Bennett DJ. Persistent inward currents in motoneuron dendrites: Implications for motor output. Muscle Nerve. 2005;31(2):135–156. doi:10.1002/mus.20261

14. Heckman CJ, Mottram C, Quinlan K, Theiss R, Schuster J. Motoneuron excitability: the importance of neuromodulatory inputs. Clin Neurophysiol Off J Int Fed Clin Neurophysiol. 2009;120(12):2040–2054. doi:10.1016/j.clinph.2009.08.009

15. Gorassini M, Yang JF, Siu M, Bennett DJ. Intrinsic Activation of Human Motoneurons: Possible Contribution to Motor Unit Excitation. J Neurophysiol. 2002;87(4):1850–1858. doi:10.1152/jn.00024.2001

16. Goodlich BI, Del Vecchio A, Horan SA, Kavanagh JJ. Blockade of 5-HT2 receptors suppresses motor unit firing and estimates of persistent inward currents during voluntary muscle contraction in humans. J Physiol. 2023;601(6):1121–1138. doi:10.1113/JP284164

17. Afsharipour B, Manzur N, Duchcherer J, et al. Estimation of self-sustained activity produced by persistent inward currents using firing rate profiles of multiple motor units in humans. J Neurophysiol. 2020;124(1):63–85. doi:10.1152/jn.00194.2020

18. Oliveira DS de, Carbonaro M, Raiteri BJ, Botter A, Ponfick M, Del Vecchio A. The discharge characteristics of motor units innervating functionally paralyzed muscles. J Neurophysiol. 2025;133(2):343–357. doi:10.1152/jn.00389.2024

19. Ting JE, Del Vecchio A, Sarma D, et al. Sensing and decoding the neural drive to paralyzed muscles during attempted movements of a person with tetraplegia using a sleeve array. J Neurophysiol. 2021;126(6):2104–2118. doi:10.1152/jn.00220.2021

20. Oliveira DS de, Ponfick M, Braun DI, et al. A direct spinal cord–computer interface enables the control of the paralysed hand in spinal cord injury. Brain. 2024;147(10):3583–3595. doi:10.1093/brain/awae088

21. Sîmpetru RC, Souza De Oliveira D, Ponfick M, Vecchio AD. Identification of Spared and Proportionally Controllable Hand Motor Dimensions in Motor Complete Spinal Cord Injuries Using Latent Manifold Analysis. IEEE Trans Neural Syst Rehabil Eng. 2024;32:3741–3750. doi:10.1109/TNSRE.2024.3472063

22. Sîmpetru RC, Braun DI, Simon AU, et al. MyoGestic: EMG interfacing framework for decoding multiple spared motor dimensions in individuals with neural lesions. Sci Adv. 2025;11(15):eads9150. doi:10.1126/sciadv.ads9150

23. Braun DI, Oliveira DS de, Bayer P, Ponfick M, Kinfe TM, Vecchio AD. NeurOne: High-performance Motor Unit-Computer Interface for the Paralyzed. medRxiv. Preprint posted online September 26, 2023:2023.09.25.23295902. doi:10.1101/2023.09.25.23295902

24. Sîmpetru RC, Arkudas A, Braun DI, et al. Learning a hand model from dynamic movements using high-density EMG and convolutional neural networks. IEEE Trans Biomed Eng. 2024;71(12):3556–3568. doi:10.1109/TBME.2024.3432800

25. Holobar A, Zazula D. Multichannel Blind Source Separation Using Convolution Kernel Compensation. IEEE Trans Signal Process. 2007;55(9):4487–4496. doi:10.1109/TSP.2007.896108

26. Del Vecchio A, Holobar A, Falla D, Felici F, Enoka RM, Farina D. Tutorial: Analysis of motor unit discharge characteristics from high-density surface EMG signals. J Electromyogr Kinesiol. 2020;53:102426. doi:10.1016/j.jelekin.2020.102426

27. McGill KC, Lateva ZC, Marateb HR. EMGLAB: An interactive EMG decomposition program. J Neurosci Methods. 2005;149(2):121–133. doi:10.1016/j.jneumeth.2005.05.015

28. Tomczak M, Tomczak E. The need to report effect size estimates revisited. An overview of some recommended measures of effect size. Published online 2014.

29. Cohen J. Statistical Power Analysis for the Behavioral Sciences. 2. ed., reprint. Psychology Press; 2009.

30. Bao S, Lei Y. Motor unit activity and synaptic inputs to motoneurons in the caudal part of the injured spinal cord. J Neurophysiol. 2024;131(2):187–197. doi:10.1152/jn.00178.2023

31. Cakici AL, Osswald M, Souza de Oliveira D, et al. A generalized framework for the study of spinal motor neurons controlling the human hand during dynamic movements. In: Proceedings of the 2022 44th Annual International Conference of the IEEE Engineering in Medicine & Biology Society (EMBC). 2022:4115–4118. doi:10.1109/EMBC48229.2022.9870914

32. Oßwald M, Cakici AL, Souza De Oliveira D, Braun DI, Farina D, Del Vecchio A. Task-specific motor units in the extrinsic hand muscles control single-and multidigit tasks of the human hand. J Appl Physiol. 2025;138(5):1187–1200. doi:10.1152/japplphysiol.00911.2024

33. Johnson MD, Heckman CJ. Gain control mechanisms in spinal motoneurons. Front Neural Circuits. 2014;8. doi:10.3389/fncir.2014.00081

34. Murray M, Goldberger ME. Restitution of function and collateral sprouting in the cat spinal cord: The partially hemisected animal. J Comp Neurol. 1974;158(1):19–36. doi:10.1002/cne.901580103

35. Fouad K, Pedersen V, Schwab ME, Brösamle C. Cervical sprouting of corticospinal fibers after thoracic spinal cord injury accompanies shifts in evoked motor responses. Curr Biol. 2001;11(22):1766–1770. doi:10.1016/S0960-9822(01)00535-8

36. Oudega M, Perez MA. Corticospinal reorganization after spinal cord injury. J Physiol. 2012;590(16):3647–3663. doi:10.1113/jphysiol.2012.233189

37. Bareyre FM, Kerschensteiner M, Raineteau O, Mettenleiter TC, Weinmann O, Schwab ME. The injured spinal cord spontaneously forms a new intraspinal circuit in adult rats. Nat Neurosci. 2004;7(3):269–277. doi:10.1038/nn1195

38. Walker JR, Detloff MR. Plasticity in Cervical Motor Circuits following Spinal Cord Injury and Rehabilitation. Biology. 2021;10(10). doi:10.3390/biology10100976

