## Supplemental Materials for "Physiological and pathological motor unit phenotypes coexist within single motor pools after cervical SCI"

### Supplementary Materials

#### Supplementary Methods

##### Exclusion Criteria

Participants were excluded if any of the following applied:

1. Age below 18 or above 80 years
2. Pregnancy
3. Use of analgesic or anticoagulant medication and/or alcohol within 24 hours prior to participation
4. Drug addiction, defined as use of illegal substances or use of legal substances above permitted limits
5. Pre-existing neurological or psychiatric illness
6. Postoperative complications
7. Inability to cooperate with experimental instructions
8. Poor skin condition due to pressure sores or electrode-related irritation in the recording region
9. History of poorly controlled epileptic seizures
10. Use of a cardiac pacemaker
11. Active cancerous tumor in the recording or stimulation region
12. Exposed orthopedic metal in the recording region
13. Unhealed bone fracture in the recording region
14. Botulinum toxin administration in the recording region
15. Pre-existing musculoskeletal illness (waived for participants in the open-loop sEMG protocol)

#### Detailed iEMG Implantation Protocol

Implant site identification presented challenges due to advanced muscle atrophy in chronic SCI and the high selectivity of fine-wire electrodes. Participants were seated in their wheelchair (P1, P3) or reclining in a hospital bed (P2), the latter to accommodate the longer preparation time required for electrode implantation. Maximum voluntary contractions were obtained for each digit (three attempts, 3 s each, 30 s rest between attempts) to establish proportional control scaling. Offsets were calculated by comparing resting baseline activity to account for the influence of tonic MU activity on baseline signal levels. An ArtUs EXT-2H ultrasound system (15-MHz linear transducer, L15-7H40-A5; Telemed, Vilnius, Lithuania) was used to identify target muscles. The associated digit was passively manipulated to confirm muscle identity under ultrasound visualization.

Electrodes were implanted under real-time ultrasound guidance, with the cannula tip visible as a hyperechoic point. Electrode position was confirmed by observing cannula movement during passive digit manipulation. If positioning was suboptimal, insertion depth was adjusted. The cannula was then withdrawn, leaving fine wires in situ. Failed implantations (electrode pullout during cannula retraction or inability to reach target) required repeat attempts. Successfully implanted electrodes were connected to the amplifier via custom four-channel bipolar adapters with spring-loaded wire hooks.

#### Manual Editing in the Presence of Tonic Activity

Manual editing followed established procedures, with iterative inspection of each unit's discharge pattern, action potential waveform, and inter-spike interval statistics. The MU filter was re-applied after removing ambiguous spikes to recover missed discharges. In this population, tonic units presented a specific editing challenge: because firing persisted through rest epochs at stable rates, the conventional visual cue of a gap in discharge used to identify decomposition errors was absent. Editing decisions for these units therefore relied primarily on waveform consistency across active and rest epochs and on stability of the inter-spike interval distribution, rather than on the presence of silent periods.

#### MUAP Tracking

To assess whether cross-session discordance reflected genuine behavioral change or threshold sensitivity, a continuous derecruitment score was computed for each MU as a weighted combination of silent fraction and normalized derecruitment delay (0.7 × (1 - silent fraction) + 0.3 × normalized delay), mapping each unit to a single value on a [0, 1] axis where 0 corresponds to fully controllable behavior and 1 to fully tonic behavior. For each matched pair, the absolute difference in derecruitment scores between sessions was computed. A permutation test (10,000 iterations) compared the observed mean absolute difference for matched pairs against a null distribution generated by randomly reassigning session labels, testing whether matched units exhibited more similar derecruitment behavior than expected by chance.

*Supplemental Figure 1: Cross-session motor unit tracking confirms phenotype stability.*

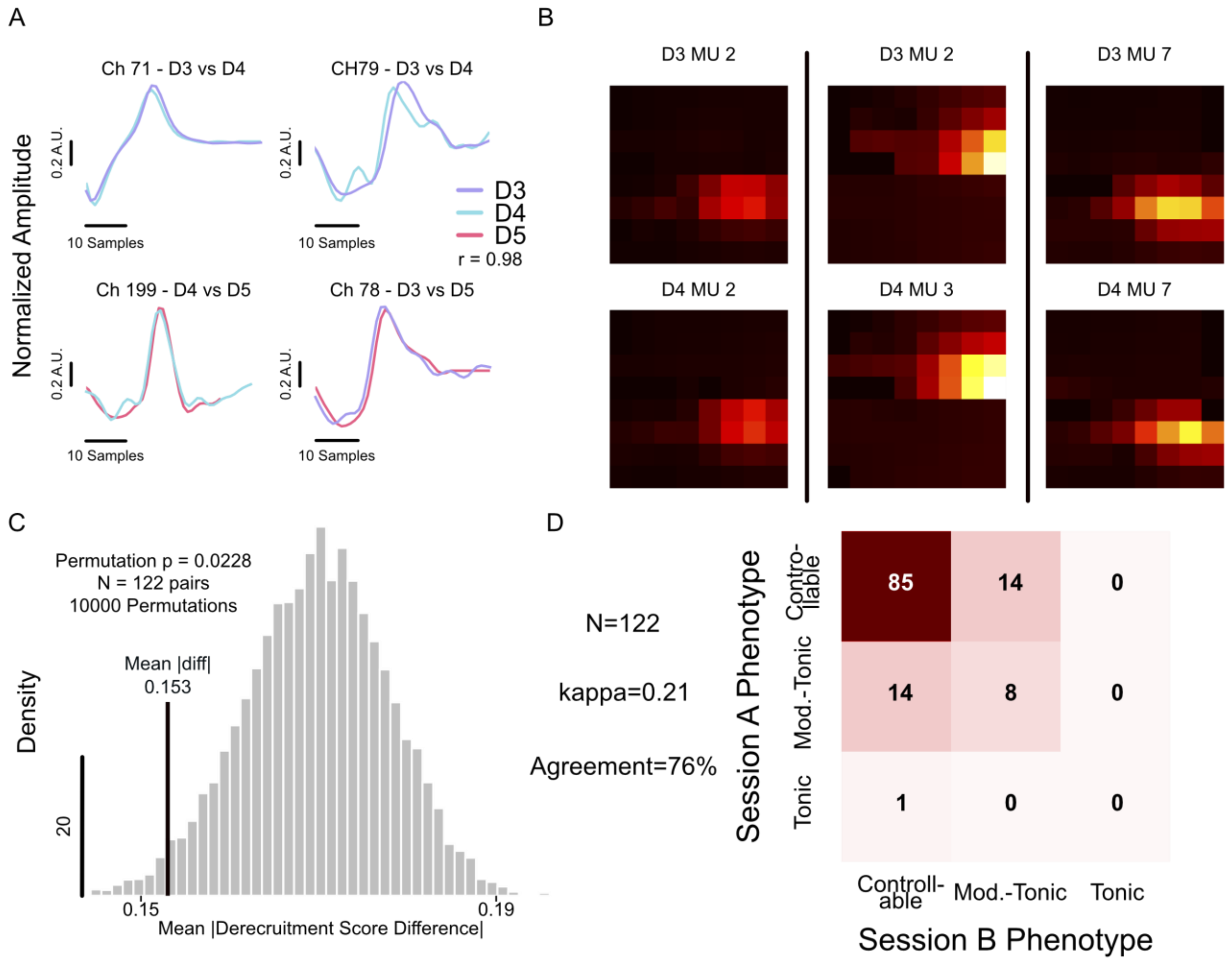


*Supplemental Figure 1 - Cross-session motor unit tracking confirms phenotype stability.*

***(A)*** *Spike-triggered average (STA) waveforms for a representative matched MU recorded across three sessions (D3, D4, D5) of the closed-loop protocol in P1. Each subplot shows the normalized STA waveform at the same electrode channel across sessions, demonstrating consistent MUAP morphology despite electrode grid repositioning between recording days (no repositioning protocol was used). Waveform similarity was quantified by normalized 2D cross-correlation of peak-to-peak amplitude maps across the 8×8 electrode grid (r = 0.98 for the pair shown).* ***(B)*** *Spatial MUAP fingerprints for three matched MU pairs in P1, shown at decreasing similarity tiers. Each column shows one pair: top row is the peak-to-peak amplitude map from the earlier session, bottom row is the matched unit from the later session. Matching was performed using shift-tolerant normalized 2D cross-correlation with one-to-one assignment enforced via the Hungarian algorithm, restricted to same-grid comparisons (flexor vs. flexor, extensor vs. extensor).* ***(C)*** *Permutation test for continuous derecruitment stability. A continuous derecruitment score (see Supplementary Methods) was computed for each MU, mapping behavior to a single [0, 1] axis from fully controllable (0) to fully tonic (1). The grey histogram shows the null distribution of mean absolute derecruitment score differences generated by randomly reassigning session labels (10,000 permutations). The black line indicates the observed mean absolute difference for matched pairs (0.153), which was significantly smaller than expected by chance (p = 0.023, one-sided), confirming that matched units exhibit more similar derecruitment behavior across sessions than randomly paired units.* ***(D)*** *Phenotype concordance matrix for P1 (122 matched pairs across 5 closed-loop sessions). Categorical agreement was 76%. All 28 discordant pairs were controllable/modulated-tonic boundary cases; no tonic unit was reclassified as controllable or vice versa across sessions. Cohen's kappa (0.21) is deflated by the high prevalence of controllable units (~80%), which inflates chance agreement; the systematic restriction of disagreements to the controllable/modulated-tonic boundary is more informative than the kappa value alone.*

#### Temporal Filter Threshold Rationale

The 7.5–8.5 Hz rate band is centered on 8 Hz, the characteristic self-sustained firing frequency associated with sodium-mediated persistent inward currents in human motoneurons. The narrow band was chosen to maximize specificity for PIC-driven tonic firing; widening the window would increase sensitivity at the cost of flagging low-modulation voluntary contractions, which can overlap with this rate range. The windowed CV ISI threshold of 0.12 reflects the locally regular discharge expected during a self-sustained PIC plateau and is intentionally more stringent than the whole-recording CV ISI abstain range (0.10–0.30) used for phenotype classification, which aggregates across both active and rest epochs. The temporal filter was implemented as a proof-of-concept to demonstrate the feasibility of spike-train-level separation of pathological from voluntary activity; systematic optimization of these thresholds across a larger cohort is left for future work.

#### Effect size interpretation

Effect sizes were quantified using epsilon-squared ($\varepsilon^2$), calculated as H/(n−1) where H is the Kruskal-Wallis statistic [28]. Following conventions for eta-squared in ANOVA (Cohen, 1988), we apply thresholds of 0.01 (small), 0.06 (medium), and 0.14 (large). No consensus exists for epsilon-squared interpretation specifically; however, the observed phenotype effects ($\varepsilon^2$ = 0.43–0.75) substantially exceed both conventional large-effect thresholds and movement type effects ($\varepsilon^2 \leq$ 0.056), providing robust evidence that phenotype classification captures the dominant source of variance in MU derecruitment behavior.

#### Supplementary Table 1: Participant Demographics

| ID | Age | Sex | Injury Level | AIS Grade | Time Since Injury (years) | MAS Score |
|---|---|---|---|---|---|---|
| P1 | 58 | M | C8 | D | 0.5 | 2 |
| P2 | 38 | M | C5 | B | 12 | 1 |
| P3 | 61 | M | C5 | A | 10 | 2 |
| P4 | 39 | F | C5 | A | 24 | 0 |

#### Supplementary Table 2: Motor Unit Phenotype Classification Thresholds

| Criterion | Controllable Vote | Tonic Vote | Abstain |
|---|---|---|---|
| CV ISI (whole-recording)[1] | > 0.35 | < 0.25 | 0.25–0.35 |
| Fraction of silent epochs | > 0.70 | < 0.30 | 0.30–0.70 |
| Mean derecruitment delay | < 0.1 s | > 0.5 s | 0.1–0.5 s |
| Per-epoch tonic fraction | < 10% | ≥ 20% | 10–20% |

**Hard constraints:**

- Derecruitment delay ≥ 0.5 s precludes controllable classification
- Derecruitment delay ≤ 0.1 s precludes tonic classification

**Classification rules:**

- Tonic: ≥3 tonic votes, OR ≥2 tonic votes with no controllable votes
- Controllable: ≥3 controllable votes, OR ≥2 controllable votes with no tonic votes
- Modulated-tonic: All other units

**Confidence adjustments:**

- Reduced by 30% if fewer than 3 rest epochs
- Reduced by 50% if CV ISI or silent fraction undefined

[1] CV ISI for classification was computed across all inter-spike intervals in the full recording (active and rest epochs combined). Because this aggregate metric reflects both voluntary and tonic firing segments, units with mixed behavior typically yield intermediate values that fall in the abstain range; classification of such units therefore depends on the remaining three criteria.

#### Supplementary Table 3. Data provenance by participant and recording modality.

| Participant | Modality | Prior Publication | Novel in this study |
|---|---|---|---|
| P1 | Closed-Loop sEMG | [18,20,23] | Phenotype classification, survival analysis, temporal filtering |
| P1 | Open-Loop sEMG | [18,20–22] [2] | Phenotype classification |
| P1 | Open-Loop iEMG | New Data | All |
| P2 | Closed-Loop sEMG | [18,20,23] | Phenotype classification, survival analysis, temporal filtering |
| P2 | Open-Loop sEMG | [18,20–22] [2] | Phenotype classification |
| P2 | Open-Loop iEMG | New Data | All |
| P3 | Open-Loop iEMG | New Data | All |
| P4 | Open-Loop sEMG | [18,20–22] [2] | Phenotype classification |

#### Supplementary Results and Analyses

##### Movement type effects:

Movement type exerted statistically significant but small effects on derecruitment delay, fraction of silent epochs, and discharge variability (all $\varepsilon^2 \leq 0.056$), with no significant effect on mean firing rate. These effects were an order of magnitude smaller than phenotype effects and did not persist after accounting for subject-level variability.

[2] Sîmpetru et al. (2024, 2025) used raw EMG data from this dataset but did not use the decomposed spike trains analyzed in the present work.

#### Supplementary Table 4. Motor unit counts by participant, recording modality, and phenotype

| Participant | Modality | Controllable | Mod.-tonic | Tonic | Total |
|---|---|---|---|---|---|
| P1 | Closed-Loop sEMG | 86 | 15 | 1 | 102 |
| P1 | Open-Loop sEMG | 2 | 5 | 3 | 10 |
| P1 | Open-Loop iEMG | 8 | 0 | 0 | 8 |
| P2 | Closed-Loop sEMG | 40 | 32 | 10 | 82 |
| P2 | Open-Loop sEMG | 6 | 27 | 16 | 49 |
| P2 | Open-Loop iEMG | 2 | 5 | 4 | 11 |
| P3 | Open-Loop iEMG | 36 | 86 | 12 | 134 |
| P4 | Open-Loop sEMG | 5 | 5 | 3 | 13 |

All motor units extracted from index flexion, two-finger pinch, and fist closure/opening movements across all recording modalities.

#### Supplementary Table 5: Phenotype comparisons (Kruskal-Wallis with Dunn's post-hoc)

| Metric | $\varepsilon^2$ | Interpretation |
|---|---|---|
| Derecruitment delay | 0.746 | Large |
| Fraction silent epochs | 0.541 | Large |
| CV ISI | 0.431 | Large |
| Mean firing rate | 0.106 | Medium |

All pairwise comparisons (controllable vs. modulated-tonic, controllable vs. tonic, modulated-tonic vs. tonic) were significant for all four metrics (all $P < 0.001$, Bonferroni-corrected).
